# Genetic profiling and *in silico* sequence analysis of *CSN2* (β-casein) and *CSN3* (κ-casein) genes in the indian buffalo (*Bubalus bubalis*)

**DOI:** 10.1101/2023.05.24.542222

**Authors:** Vinay Kumar Mehra, Satish Kumar

## Abstract

Buffalo rank second for milk production in the world and play important role in Indian economy. There are four types of caseins α-S1-casein, α-S2-casein, β-casein and κ-casein in milk. The major function of the casein protein in milk is to chelate colloidal calcium phosphate and serves as a major source of amino acids, calcium and phosphate. In buffalo milk all four casein proteins (αs1, αs2, β and κ) are encoded by four closely linked autosomal genes (CSN1S1, CSN1S2, CSN2 and CSN3 respectively) that are present on chromosome 7. Bovine CSN2 (β-casein) gene is 8.5 kb long and contains nine exons and CSN3 (κ-casein) is ∼ 13 kb long. The aim of the study was to characterize CSN2 and CSN3 genes, *In-silico* analysis of β-casein and κ-casein protein and evolutionary relationship with other species. Buffalo mammary gland tissue was collected from local slaughterhouse (New Delhi, India) and total RNA was isolated from Buffalo Mammary Epithelial Cells. The ORF region of CSN2 and CSN3 genes were amplified and sequenced for characterization. Physiochemical properties showed that both buffalo β-casein (Bu_CSN2) and κ-casein (Bu_CSN3) proteins are stable and hydrophobic in nature.The presence of high phosphorylated residues in both β-casein and κ-casein proteins residues suggested that they are involved in signal transduction processes, cell growth and metabolism. The N-glycosylation result showed that both proteins are less in foldable state. The presence of methylation and acetylation sites in both protein revealed that they are involved in different cellular process. The evolutionary analysis showed that both buffalo genes more closely to *Bos grunniens (yak)*.

## Introduction

Buffalo rank second for milk production in the world and play important role in Indian economy. Compare to cow’s milk, Buffalo milk contain higher total solids, fat, protein and calcium level and these will affect processing and yield of certain products. Milk proteins are best described by chemically, physically and genetically of all food proteins. There are two distinct groups of milk protein. First, the caseins, which are about 80% including α-S1-casein, α-S2-casein, β-casein, κ-casein and another is whey protein (β-lactoglobulin and α lactalbumin), which accounts for 20% of total protein in ruminant milk (Layman et al., 2018). The major function of the casein protein is to chelate colloidal calcium phosphate and serves as a major source of amino acids, calcium and phosphate for infants (Bawden et al., 1994). The milk protein compositions of different species differ greatly, both in the presence or absence of particular proteins and in their relative abundance. It has been reported that different caseins protein evolved from a common ancestor, by a series of duplications and exon shuffling events and different species also showed that there is few sequence conservation between different caseins within a species (Bawden et al., 1994). In buffalo milk all four casein proteins (αs1, αs2, β and κ) are encoded by four closely linked autosomal genes (CSN1S1, CSN1S2, CSN2 and CSN3 respectively) that are present on chromosome 7 and distributed in buffalo milk: αs1-casein 20.61%, αs2-casein 14.28 %, β-casein 53.45 % and κ-casein 11.66 % (Barłowska et al., 2012).

β-casein is the most hydrophobic of all the caseins and play a vital role in determining the surface properties of casein micelles and curd formation. Bovine CSN2 (β-casein) gene is 8.5 kb long and contains nine exons and eight introns. The first exon of CSN2 gene is about 44 bp and contains most of 5’ untranslated region. The remaining 12 bp of the untranslated region is encoded by Exon II. The exon II also encodes entire signal peptide and the first two codons of the mature β-casein protein *(Bonsinget et al., 1988)*. In cattle 12 alleles of β-casein have been identified (A1, A2, A3, B, C, D, E, F, H1, H2, I, G) and the buffalo β-casein differs from cattle by 6 amino acids at different positions (Vinesh et al., 2013). The most common forms of β-casein in cattle are A1 and A2, which differ by only one amino acid. Isoform A1 contains histidine and A2 proline at the 67 position of amino acid chain (Barłowskaet al., 2012).

The κ-casein contains major carbohydrate-free components and constitutes about 12% of casein protein. It stabilizes calcium sensitive caseins against precipitation and plays an important role in determining the size of casein micelles and help in curd and cheese production (Kishore et al., 2014). The length of κ-casein is about 13 kb, and most of the coding sequences are comprised in the exon-IV and structurally κ-casein is highly homologous to the fibrinogen gamma chain (Barłowska et al., 2012). There are 11 genetic variants of this gene have been identified in cattle. Among these, A and B are the most common (Kishore et al., 2014) and and these alleles differ by substitutions in 2 amino acids at positions 136 and 148 (alanine) (Otaviano et al., *2005*). Variant B is associated with higher fat, protein, and casein in comparison to AA or AB variants (Gangaraj et al., 2008). In case of buffalo allele B of CSN3 gene was most common (Barłowska et al., 2012). It is also reported that the allele B is associated with shorter coagulation time, thermal resistance, better curdles and micelles of different sizes, which make it preferable for cheese making (Azevedo et al., 2008). Thus the buffalo milk proteins particularly casein has high production and marketing significance. So, the present study was carried out to characterize CSN2 and CSN3 genes at molecular level and *In-silico* analysis for prediction of secondary structure and physiochemical properties of β-casein and κ-casein protein and evolutionary relationship with other species..

## MATERIALS AND METHODS

### Isolation and Culture of Buffalo Mammary Epithelial Cells

Buffalo mammary gland tissues were obtained from local slaughterhouse (New Delhi, India) for isolation of Buffalo Mammary Epithelial Cells (BuMEC). We followed essentially the same protocol used by Anand et al., 2012 for isolation of BuMEC with minor modifications.

### RNA Isolation and cDNA Synthesis

Total RNA from buffalo mammary tissues and BuMECs were prepared using TRIzol (Invitrogen, USA) according to manufacturer’s protocol. RNA integrity was assessed in 1.5% agarose gel electrophoresis by observing rRNA bands corresponding to 28S and 18S. Possible genomic DNA contamination in RNA preparation was removed by using DNA free kit (Ambion, USA) according to manufacturer’s protocol. The Purity of RNA was checked in UV spectrometer with the ratio of the OD at 260 nm and 280 nm being >1.8. cDNA was synthesized using Revert Aid First strand cDNA synthesis kit (Thermo Scientific, USA) by reverse transcription PCR. Briefly, 1 ug RNA was reverse transcribed using RevertAid M-MuLV reverse transcriptase (200 U/uL), RiboLock Rnase Inhibitor (20 U/uL), 10 mM dNTP mix (1 uL) oligo dT primers in 5X reaction buffer. The cDNA was stored at -20° C for further use.

### Primer designing and amplification

The primers for buffalo CSN2 (β-casein) and CSN3 (κ-casein) gene were designed using Primer3plus software based on the conserved sequence obtained from multiple sequence alignment analysis. These primers (CSN2F: 5’ CCATTCAGCTCCTCCTTCAC 3’, CSN2R: 5’ TGCCATATTTCCAGTCACAGTC 3’ and CSN3F: 5’ TACTGCCAAGCAAGAGCTGA 3’, CSN3R: 5’ GAAAGGAGCCGAATCTGTGA 3’) are targeted to compete ORF region of CSN2 and CSN3 genes respectively.

The Buffalo CSN2 and CSN3 genes were amplified using gene specific primers in Gene Pro Thermal Cycler TC-E-96G (BIOER). The PCR condition contain the following steps: initial denaturation for 3 min at 94° C followed by 35 cycles of 45 sec at 94° C, 45 sec at 58° C, 1 min at 72° C and final extension for 7 min at 72° C. The 25 μl reaction mixture contained 100 ng genomic DNA, 1 × Taq reaction buffer, 1 mM dNTPs, 50 unit of Taq polymerase and 100 ng of gene specific forward and reverse primer.

### Cloning and Sequencing

The purified PCR products were cloned into the pJET1.2 cloning vector (Thermo Scientific, #K1232). At this step, the PCR products with 3′-dA overhangs are blunted with a thermostable DNA blunting enzyme and then ligated to the linearized pJET1.2 cloning vector. The cloned PCR products were transformed into Top10 (*E. coli)* competent cells. This vector contains a lethal restriction enzyme gene that is disrupted by ligation of a DNA insert into the cloning site. As a result, only bacterial cells with recombinant plasmids are able to form colonies. The transformed recombinant colony was then subjected to colony PCR and the band intensities of the amplified products were checked in 1.2% agarose gel. The desire band was eluted using QIAquick Gel Extraction Kit (QIAGEN) and sent the purified DNA for sequencing (sanger sequencing) to SciGenome Lab Pvt. Ltd. (Cochin-India).

### Sequence Analysis

The obtained buffalo CSN2 (Bu_CSN2) and CSN3 (Bu_CSN3) gene sequences were submitted to NCBI and accession number MT276580 and MT276581 were received respectively. These nucleotide sequences were assessed for the homology against the publicly available database in NCBI BLASTN (https://www.ncbi.nlm.nih.gov/BLAST) and Nucleotides sequences were then aligned with β-casein and κ-casein gene sequences of different species by using Bioedit sequence alignment Software (version 7.2.5) respectively. After sequence analysis, the buffalo Bu_CSN1S1 gene sequences was translated in to protein sequences by using Sequence Manipulation Suite (www.bioinformatics.org). Phylogenetic analyses of Bu_CSN2 and CSN3 genes were carry out by using MEGAX software (version 10.1.5) to determine the evolutionary relationship between different closely related species.

### Evolution of primary structure of buffalo α-S1-casein protein

The physicochemical properties of buffalo β-casein and κ-casein protein of Bu_CSN2 and Bu_CSN3 genes were analyzed by using ExPASy-ProtParam tool (Gasteiger et al., 2003) which computes the number of amino acids and its composition, theoretical isoelectric point (pI), molecular weight, grand average of hydropathicity (GRAVY), Instability Index, and Aliphatic Index.

### Secondary Structure Prediction

The Secondary Structures of Bu_CSN2 and Bu_CSN3 genes encoded β-casein and κ-casein protein were examined through SOPMA (Self-Optimized Prediction Method with Alignment) server (*C.Geourjon and G.Deleage, 1995*) which computes the percentage of α-helices, β turn and β-sheet.

### Prediction of Phosphorylation and Glycosylation sites

Different phosphorylation sites of buffalo β-casein and κ-casein were predicted by using NetPhos 3.1 Server tool (Blom et al., 1999). The glycosylation sites of these two proteins were determined through NetNGlyc 1.0 Server (Blom et al., 2004).

### Prediction of Methylation and Acetylation sites

The potential methylation and acetylation sites of Bu_CSN2 and Bu_CSN3 genes encoded αs1 β-casein and κ-casein protein were predicted by using an *in-silico* tool PLMLA (Prediction of lysine methylation and lysine acetylation) respectively (Shi et al., 2012).

### Prediction of Motifs

The presences of motifs in buffalo β-casein and κ-casein proteins were analyzed through MEME (Multiple Extraction-Maximization for Motif Elicitation) tool (Bailey et al., 2009).

## Result and Discussion

### Amplification of buffalo CSN1S1 gene and sequence analysis

The ORF region of buffalo CSN2 and CSN3 genes were amplified and resulted product size of 795 bp and 745 bp respectively (fig.1). The sequence analysis through Bioedit sequence alignment programme showed that there were no nucleotide changes in the coding sequence of Bu_CSN2 and Bu_CSN3 gene (Fig.2 & Fig.3)

**Fig. 1.**
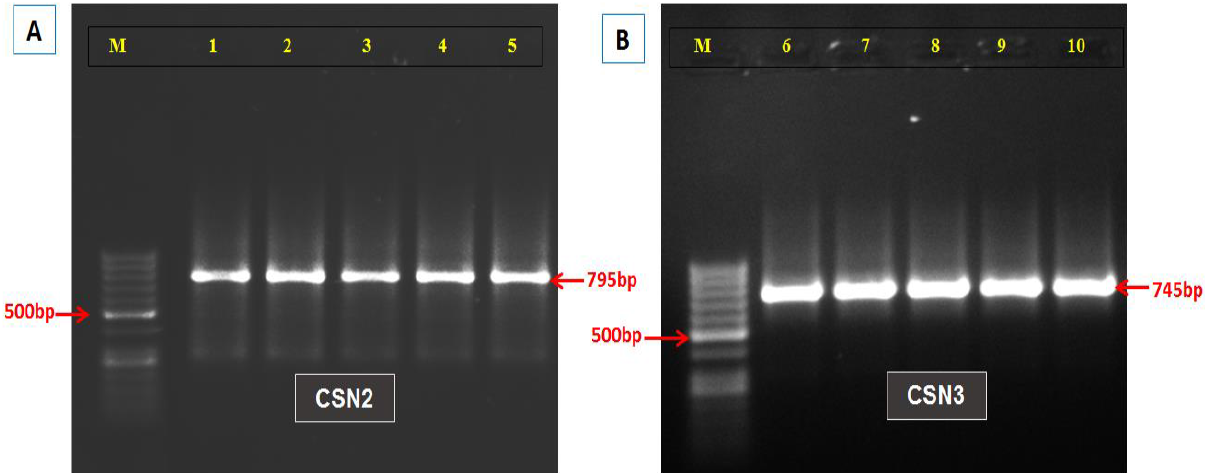
Gel electrophoresis of (A) Bu_CSN2 and (B) BU_CSN3 cloned cells PCR products. Lane 1-5 CSN2 gene, Lane 6-10 CSN3 gene and M-50 bp ladder (5 samples for each gene)

**Fig. 2.**
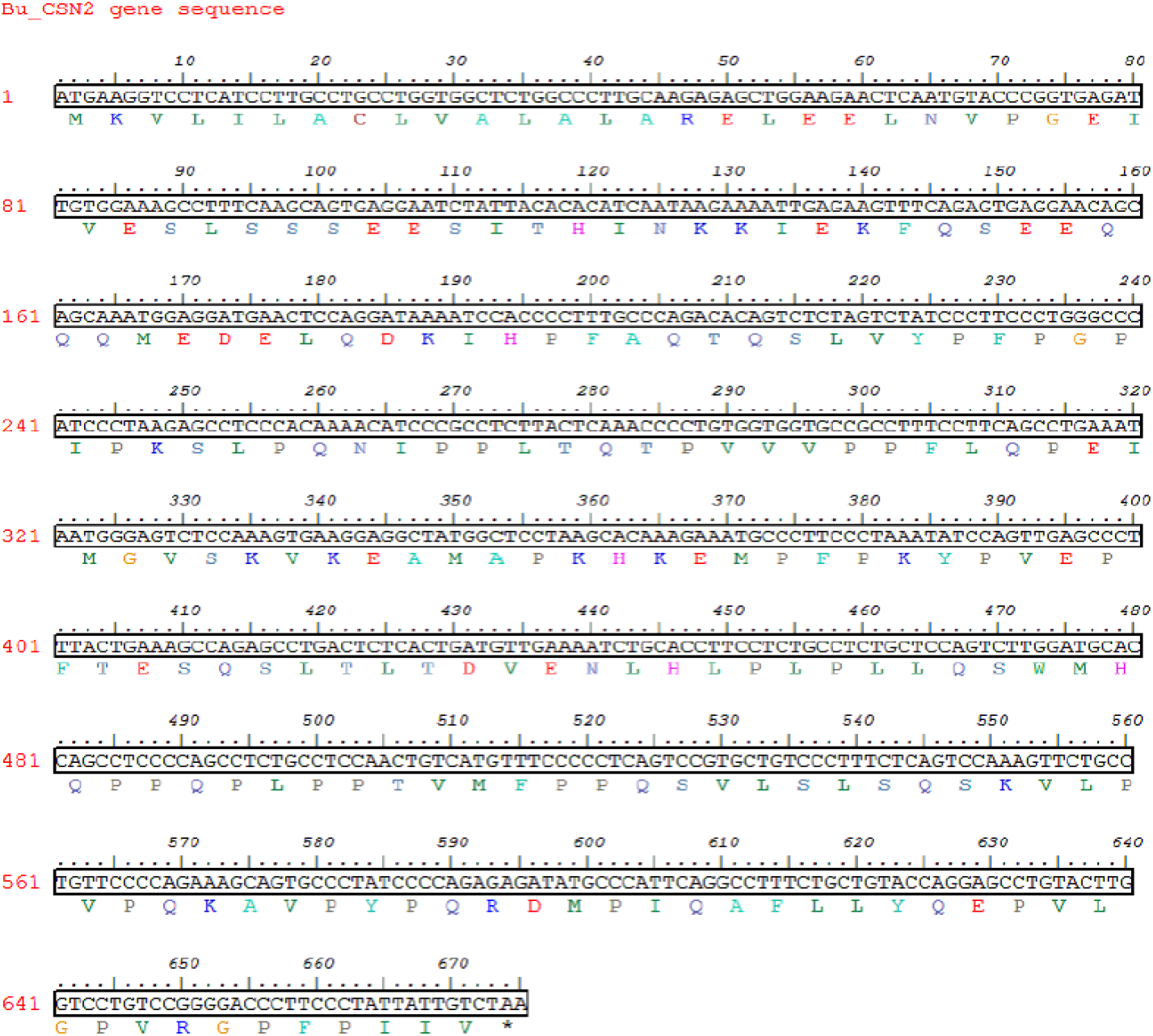
Complete cDNA sequence (in box) of B*ubalus bulalis* CSN2 gene and corresponding protein sequence is reported in color, whereas asterisk represents the termination stop codon

**Fig. 3.**
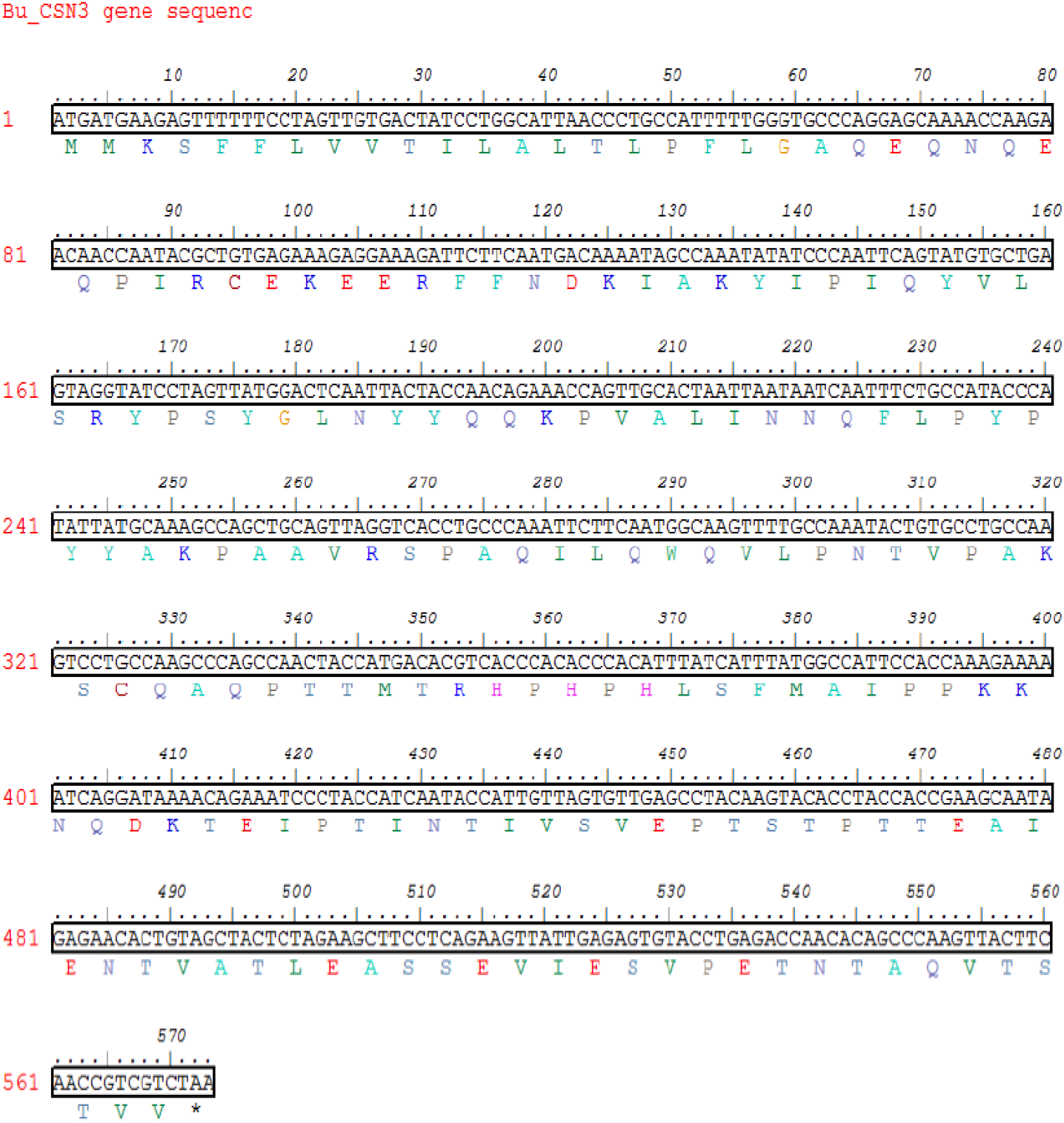
Complete cDNA sequence (in box) of B*ubalus bulalis* CSN3 gene and corresponding protein sequence is reported in color, whereas asterisk represents the termination stop codon

### Sequence Homology

The obtained nucleotide and translated amino acid sequences of Bu_CSN2 and Bu_CSN3 gene were analyzed for the sequence homology through NCBI BLASTN and BLASTP tools respectively. The nucleotide sequence of Bu_CSN2 showed 100% identity along with 100% query cover with *Bubalus bubalis* haplotype 2 casein beta (CSN2) mRNA, complete cds. The translated amino acid sequence of Bu_CSN2 gene also showed higher level of similarities with beta-casein (*Bubalus bubalis*) protein sequence. Similarly Bu_CSN3 nucleotide sequence showed 100% similarities with *Bubalus bubalis* casein kappa (CSN3), mRNA.

### Evolution of Primary Structure

The primary structures of Bu_CSN2 and Bu_CSN3 gene encoded β-casein and κ-casein protein sequences were evaluated by some parameters such as the number of amino acids, pI, molecular weight, Instability Index, Aliphatic Index and Grand average of hydropathicity (GRAVY) respectively (Table 1). The high pI value represents the basic nature of amino acid and low pI indicates acidic amino acids. The result of ExPASy-ProtParam showed that both buffalo β-casein (Bu_CSN2) and κ-casein (Bu_CSN3) proteins are of acidic in nature. The stability of a protein is determined by the instability index; the value < 40 is predicted to be stable and > 40 is unstable. In this context, our data revealed that both the proteins are not stable (Table 1). The Aliphatic Index of a protein is based on the presence of aliphatic amino acids (alanine, valine, isoleucine and leucine) residing in the aliphatic side chain of that protein. Higher value of aliphatic index indicates the more thermotolerant protein (Ertuğrul Filiz and Ibrahim Koc 2014). In this study, both Bu_CSN2 and Bu_CSN2 proteins contain comparatively high percentage of aliphatic amino acids showing the thermostable nature of these proteins. The negative GRAVY value (Table 1) for Bu_CSN2 and Bu_CSN3 protein s indicated the hydrophobic nature of these proteins (Ertugrul Filiz and Ibrahim Koc 2014).

**Table 1.** Physicochemical characteristics of Bu_CSN2 (β-casein) and Bu_CSN3 (κ-casein) proteins.

| Physicochemical parameters | Bu_CSN2<br>(β-casein) | Bu_CSN3<br>(κ-casein) |
| --- | --- | --- |
| Number of amino acids | 224 | 190 |
| Theoretical pI | 5.26 | 6.83 |
| Molecular weight | 25106.44 | 21397.56 |
| Instability index | 97.80 | 49.60 |
| Aliphatic index | 97.81 | 84.16 |
| Grand average of hydropathicity (GRAVY) | -0.130 | -0.259 |
| Total number of negatively charged residues (Asp + Glu) | 23 | 15 |
| Total number of positively charged residues (Arg + Lys) | 16 | 15 |

### Evaluation of Secondary Structure

The secondary structures of buffalo β-casein and κ-casein proteins were predicted by SOPMA server and showed in Fig. 4. The protein flexibility is defined by the presence of random coils. High percentage 60.71% and 66.84% of random coils in β-casein (Bu_CSN2) and κ-casein (Bu_CSN3) proteins confirmed that both are flexible structures respectively (Mark V. Berjanskii and David S. Wishart 2008). Similarly, low percentage of alpha helix in β-casein (33.48%) and κ-casein (17.89%) proteins indicated that these proteins are not thermostable, since thermophilic proteins have abundance of alpha helices (sandeep et al.,2000).

**Fig. 4.**
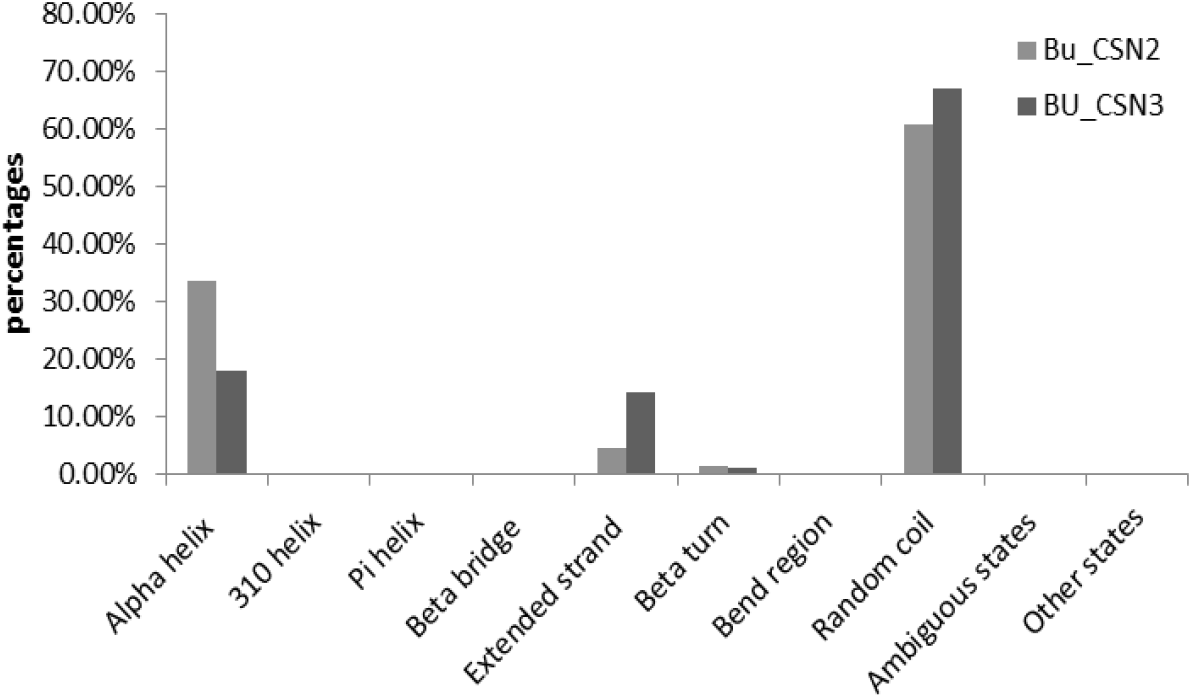
Graphical representation of the secondary structure analysis using SOPMA server of Bu_CSN2 (β-casein) and Bu_CSN3 (κ-casein) proteins.

### Evaluation of Phosphorylation and Glycosylation sites of β-casein and κ-casein protein sequences

Phosphorylations of protein change the activity of cellular proteins quickly from one state to another. Thus, protein phosphorylation is identified as a key step in various cell signaling pathways (Blom et al 2004). The insertion or deletion of phosphate group resulted in alteration of protein function. The result of NetPhos 3.1 Server of β-casein (Bu_CSN2) protein (Fig.5a) showed that there were 16 serine, 8 threonine and 4 tyrosine but only 11 serine, 6 threonine and 0 tyrosine were above threshold levels which can be predicted as phosphorylated. Similarly in Bu_CSN3 (κ-casein) protein (Fig.5b) 10 serine, 9 threonine and 1 tyrosine can be predicted as phosphorylated. So this result showed that presence of 17 and 20 phosphorylated residues in the buffalo β-casein (Bu_CSN2) and κ-casein proteins (Bu_CSN3) indicated that they are involved in signal transduction processes, cell growth and metabolism, respectively (Batra et al., 2019).

**Fig. 5.**
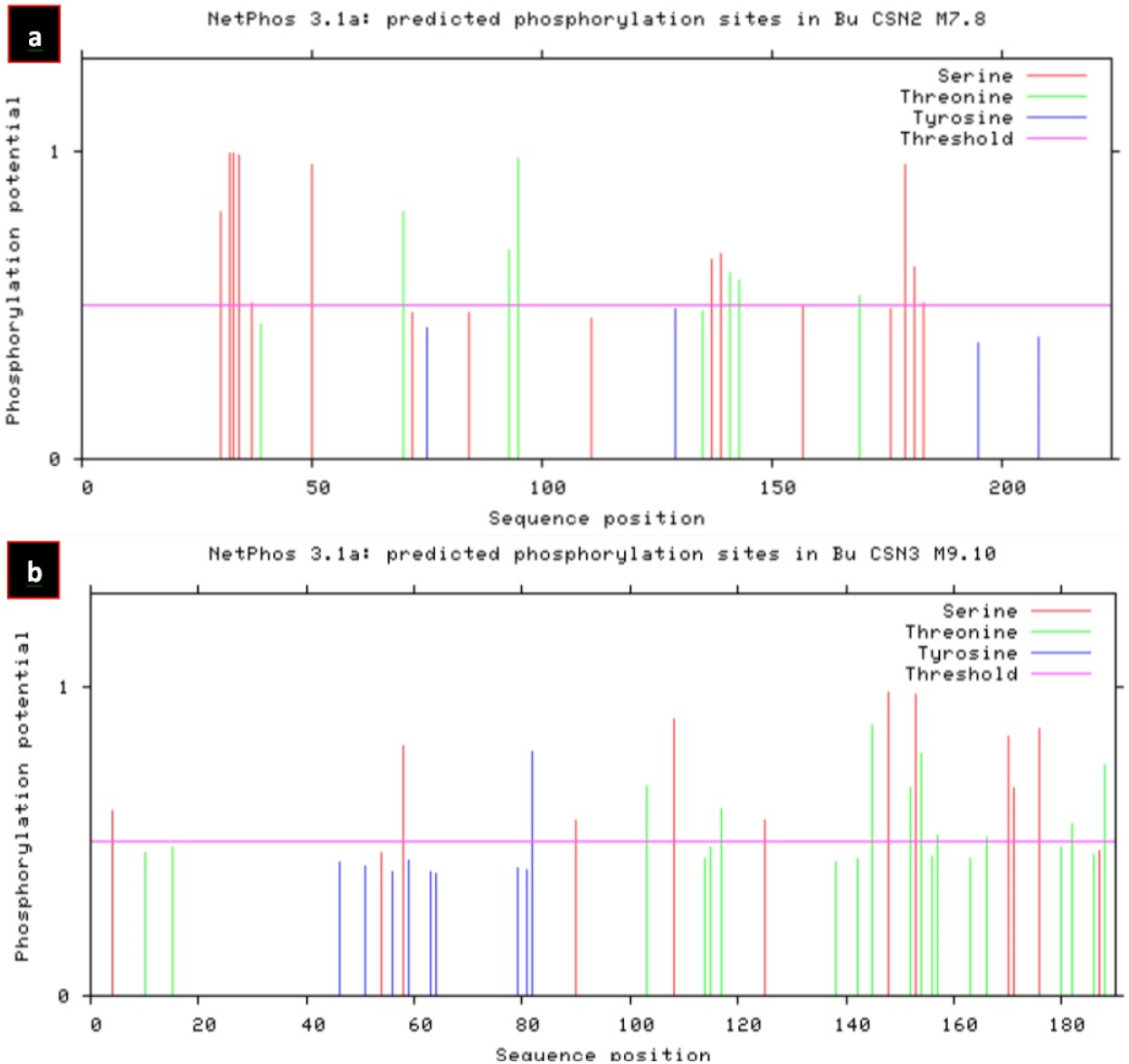
Graphically representation of Phosphorylation site predicted by NetPhos 3.1 Server (a) β-casein (Bu_CSN2) and (b) κ-casein (Bu_CSN3) protein showing different positions of Phosphorylation site.

The structure, folding and stability of a protein is also determined by the glycosylation pattern. The NetNGlyc 1.0 Server glycosylation prediction result (Fig.6a & b) showed that not a single amino acid residue was N-glycosylated in both proteins (Bu_CSN2 & Bu_CSN3), which suggested that these proteins are less stable and less in foldable state (Shental Bechor, D. & Levy Y., 2008).

**Fig. 6.**
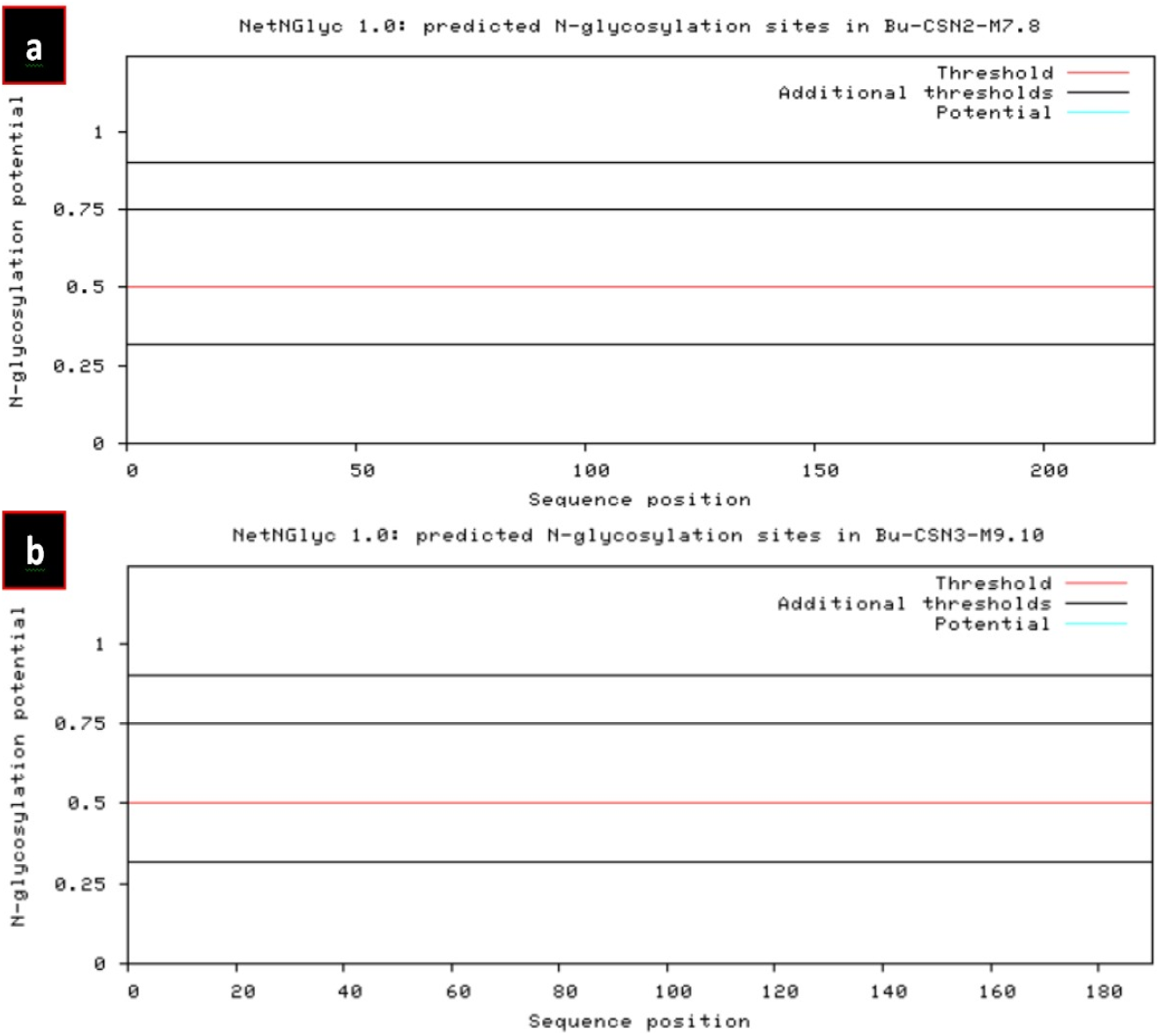
Graphically representation of N-glycosylation site predicted by NetNGlyc 1.0 Server (a) β-casein (Bu_CSN2) and (b) κ-casein (Bu_CSN3) protein

### Prediction of Methylation and Acetylation sites β-casein and κ-casein protein sequences

PLMLA programme result showed that there were 9 methylated lysine and 7 acetyl lysine sites present in β-casein (Bu_CSN2) protein sequence (Table 2). Similarly in κ-casein (Bu_CSN3) protein 4 methylated lysine and 7 acetyl lysine sites found (Table 3). These covalent modification of specific lysine residue in both the proteins suggested that they play different role in cellular process including gene expression, chromosome assembly and DNA repair etc. So predictions of methylation and acetylation sites in β-casein and κ-casein proteins are very helpful for identification of structural and functional properties of the proteins (Batra et al., 2019).

**Table 2.**
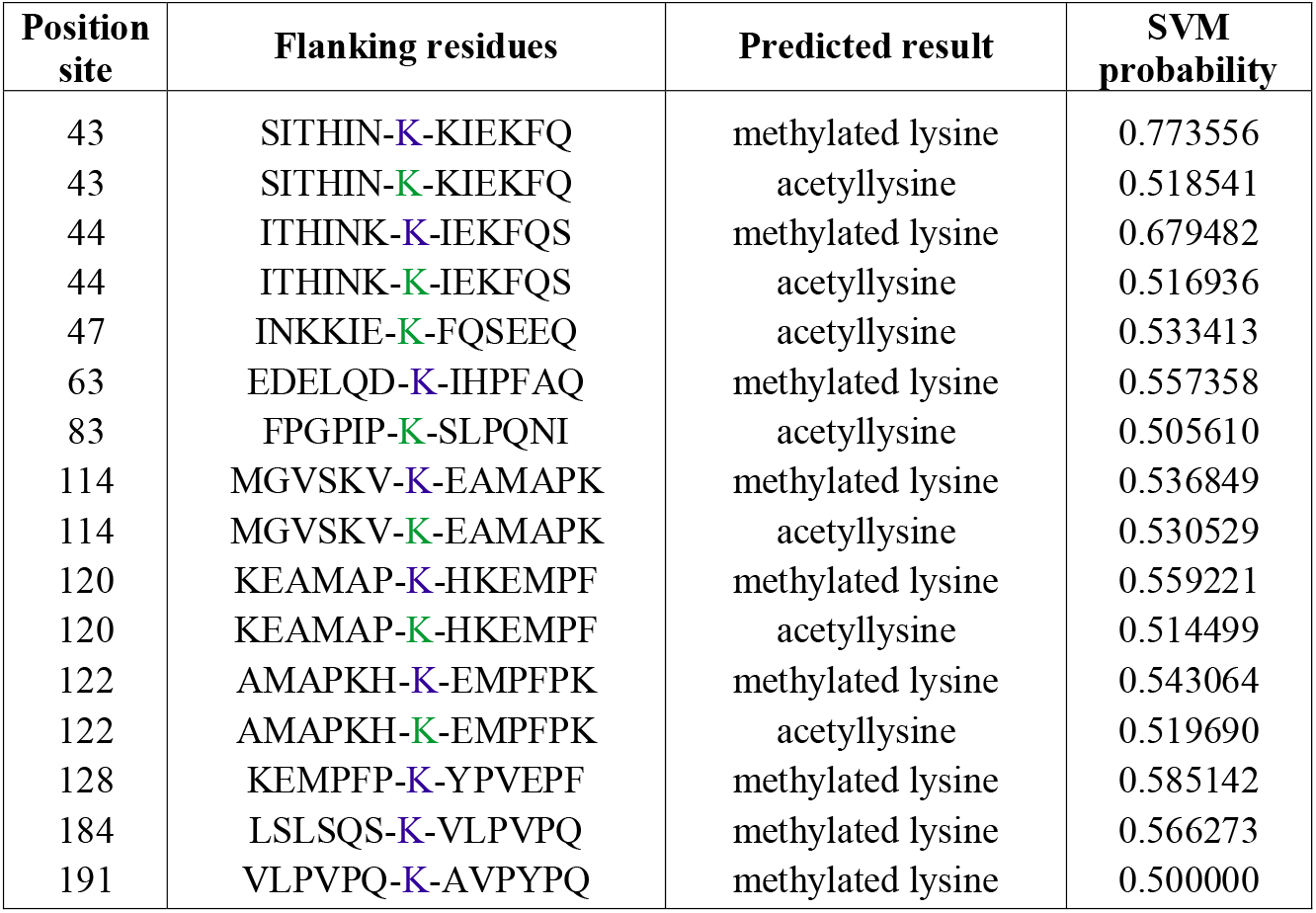
Different acetylation and methylation sites in β-casein (Bu_CSN2) protein.

**Table 3.** Different acetylation and methylation site in κ-casein (Bu_CSN3) protein.

| Position site | Flanking residues | Predicted result | SVM probability |
| --- | --- | --- | --- |
| 34 | QPIRCE-**K**-EERFFN | methylyated lysine | 0.551068 |
| 34 | QPIRCE-**K**-EERFFN | acetyllysine | 0.531975 |
| 42 | ERFFND-**K**-IAKYIP | methylyated lysine | 0.511779 |
| 42 | ERFFND-**K**-IAKYIP | acetyllysine | 0.545781 |
| 45 | FNDKIA-**K**-YIPIQY | methylyated lysine | 0.607944 |
| 45 | FNDKIA-**K**-YIPIQY | acetyllysine | 0.515690 |
| 67 | LNYYQQ-**K**-PVALIN | acetyllysine | 0.529001 |
| 84 | PYPYYA-**K**-PAAVRS | methylyated lysine | 0.523968 |
| 84 | PYPYYA-**K**-PAAVRS | acetyllysine | 0.535584 |
| 133 | MAIPPK-**K**-NQDKTE | acetyllysine | 0.510594 |
| 137 | PKKNQD-**K**-TEIPTI | acetyllysine | 0.500000 |

### Prediction of motifs in β-casein and κ-casein protein

Prediction of motifs in DNA or protein sequences enable the users to find signals (motifs). For example binding sites for the same transcription factor and protein-protein interaction domains etc (Bailey et al., 2006). MEME tool utilize statistical modeling algorithm such as probabilistic MEME and discrete models DREME fusion with GLAM2 algorithm to predict conserved motifs. This tool utilizes unaligned amino acid sequences equivalent to target protein sequence as input and creates stretches of conserved motifs as output (Chakraborty et al., 2020). In present study, the templates which have more than 90% and 85% similarities with buffalo β-casein (Bu_CSN2) and κ-casein (Bu_CSN3) proteins were selected as the input and the location of conserved motifs between these sequences are presented as the output (Fig.7 & Fig.8).

**Fig. 7.**
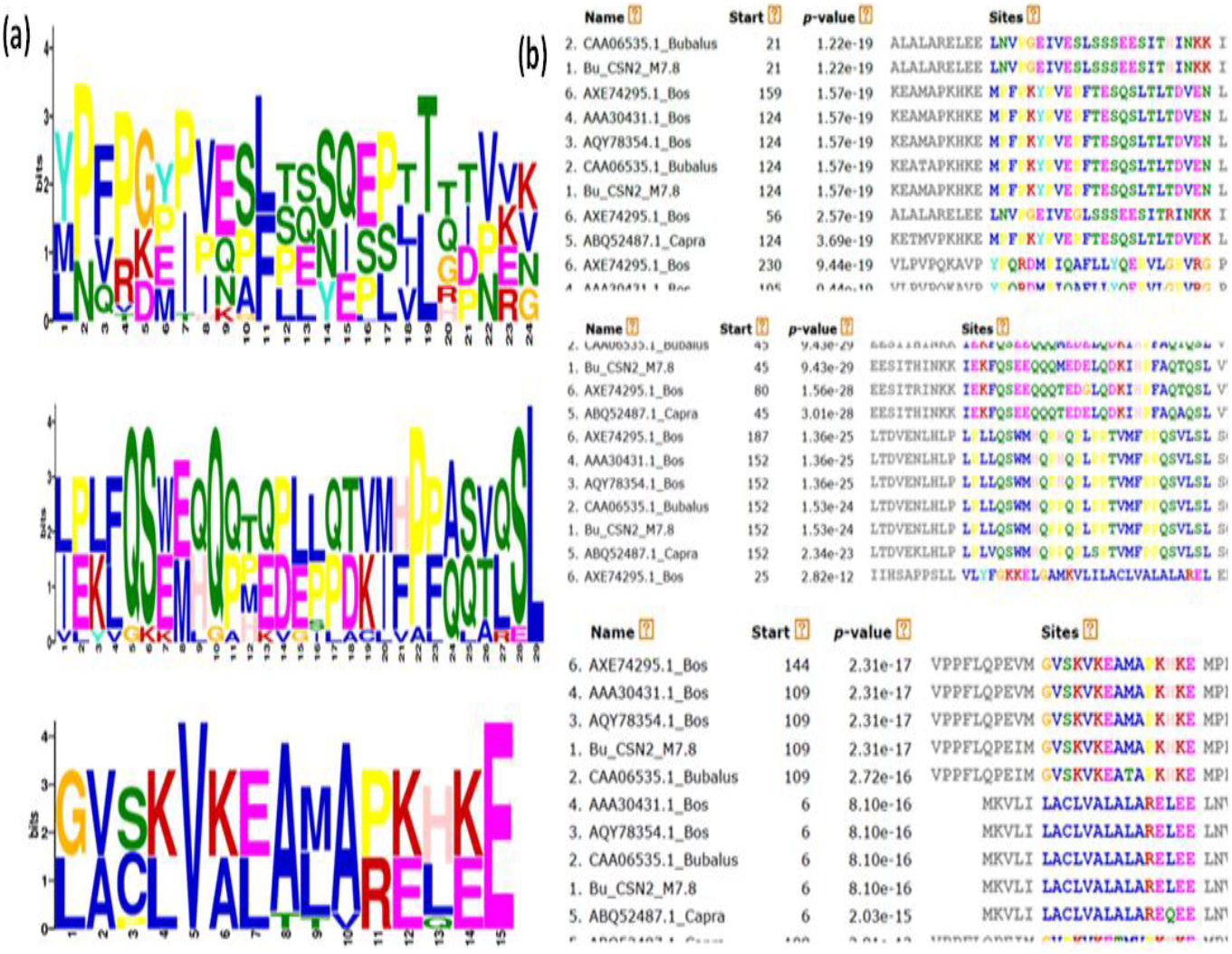
Conserved motif sequence in β-casein protein through MEME suit. (a) MEME programme showed three conserved motif region (The height of a letter shows its relative frequency at the given position in the motif), (b) templates showed more than 90% similarities with buffalo β-casein (Bu_CSN2) protein

**Fig. 8.**
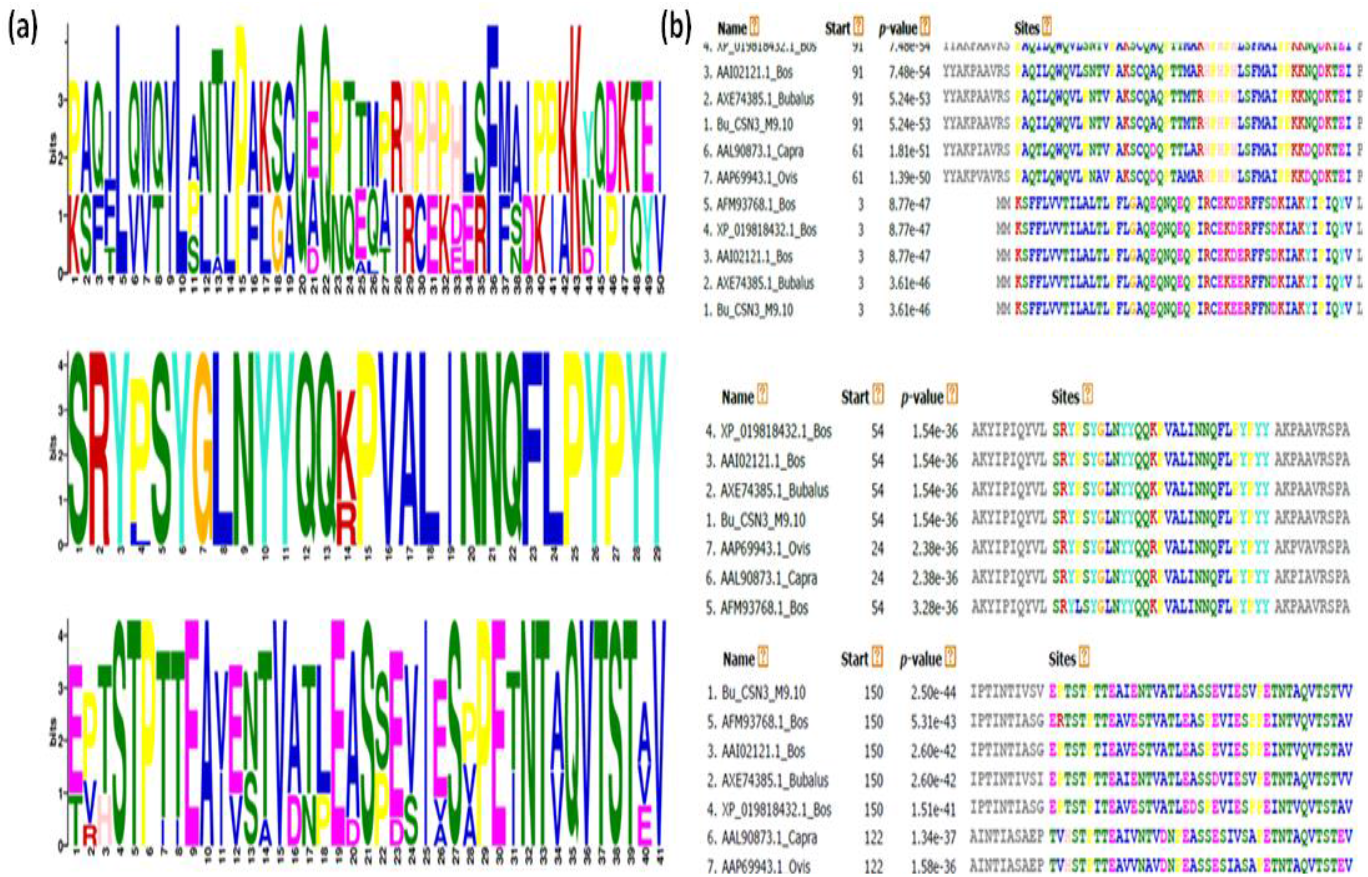
Conserved motif sequence in κ-casein protein through MEME suit. (a) MEME programme showed three conserved motif region (The height of a letter shows its relative frequency at the given position in the motif), (b) templates showed more than 85% similarities with buffalo κ-casein (Bu_CSN3) proteins protein.

### Phylogenetic analysis of CSN2 and CSN3gene sequence

Phylogenetic tree of buffalo (*bubalus bubalis*) CSN2 and CSN3 genes were constructed with the help of MEGAX (version 10.1.5) program using UPGMA method with the bootstrap values from 1000 replicates against the nucleotide sequences of the β-casein (CSN2) genes of cattle (*Bos taurus and Bos indicus)*, sheep (*Ovis aries*), goat (*Capra aegagrus hircus*), yak (*Bos grunniens*) and human (*Homo sapiens*) obtained from NCBI database respectively. In the evolutionary relationships, it has been observed that cattle, yak and buffalo formed a cluster. The buffalo CSN2 (Bu_CSN2) gene showed 97% closer relationship with *Bos grunniens (yak)* (Fig.9). On the other hand, sheep and goat formed another cluster (100%) with a closer relationship. However, human was placed as an out-group in the tree. Similarly buffalo κ-casein (Bu_CSN3) gene showed 97% similarity with κ-casein (CSN3) gene of yak (Fig.10).

**Fig. 9.**
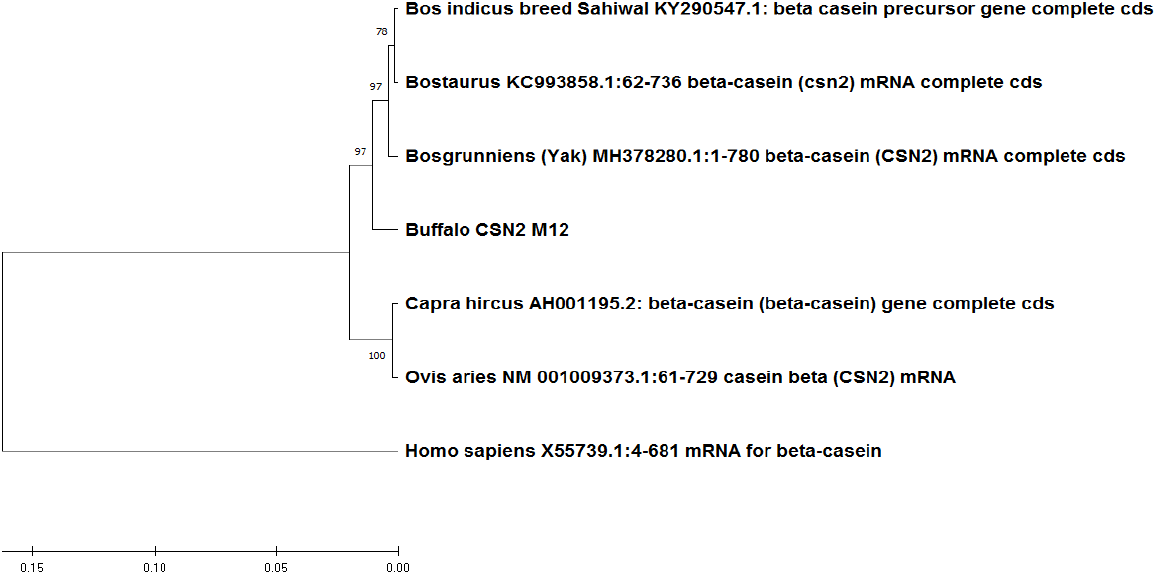
Evolutionary relationships of Bu_CSN2 gene based nucleotide sequences of the CSN2 gene of different species. The analysis was carried out with the UPGMA method with the bootstrap values from 1000 replicates, as indicated by the numerical values on the nodes.

**Fig. 10.**
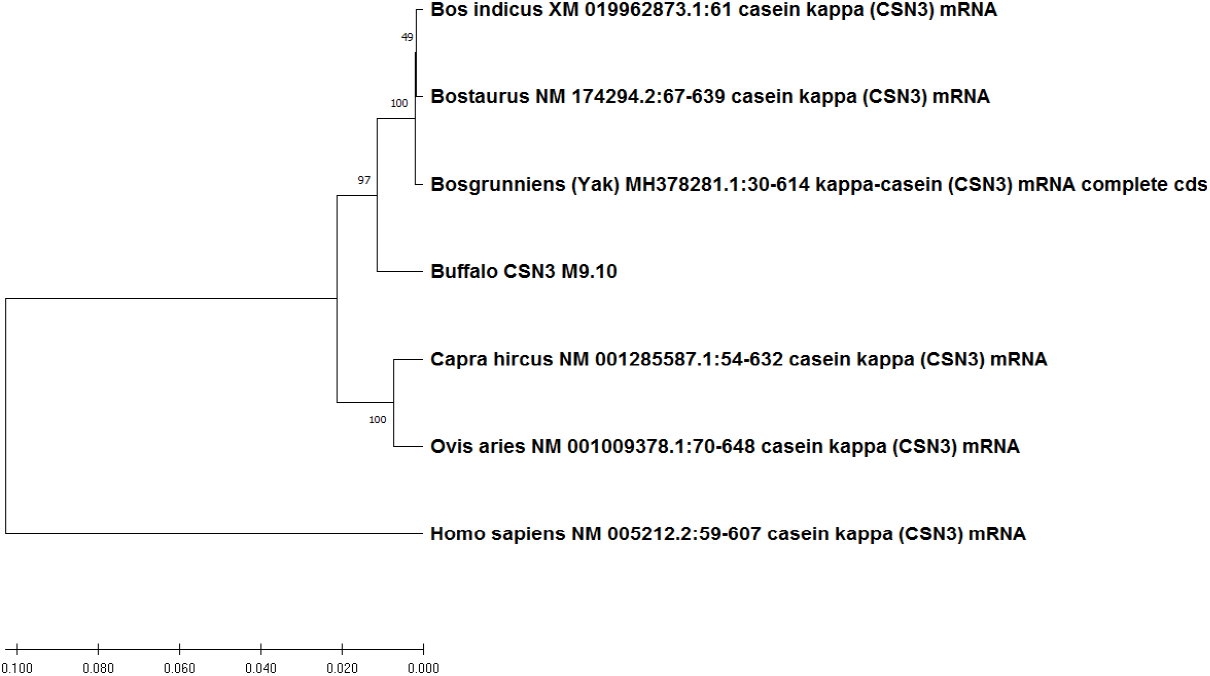
Evolutionary relationships of Bu_CSN23 gene based nucleotide sequences of the CSN3 gene of different species. The analysis was carried out with the UPGMA method with the bootstrap values from 1000 replicates, as indicated by the numerical values on the nodes.

## Conclusion

Caseins are milk proteins secreted by mammary gland epithelial cells and associated with milk quality and milk yield. β-casein protein plays a vital role in determining the surface property of casein micelle and κ-casein stabilize this structure. The complete cDNA sequences of buffalo CSN2 and CSN3 gene share similar organization with cattle, sheep, goat and yak CSN2 and CSN3 gene respectively. Prediction of protein structure allows the understanding of protein’s function. The computational analysis of buffalo β-casein and κ-casein protein structure showed that this both protein are acidic in nature and not so stable. The hydropathicity index showed that β-casein and κ-casein protein are hydrophobic in nature. It was also found that both the protein have flexible structure. Further, it was also identified that buffalo β-casein and κ-casein proteins have more phosphorylation site but no site for glycosylation. The methylation and acetylation of specific lysine residue in this proteins suggested that they may involve in various epigenetic regulation. The presence of different motifs in buffalo β-casein and κ-casein proteins indicated that they show protein-protein interactions and involved in various cellular process. From evolutionary analysis, it concluded that both buffalo gene (CSN2 and CSN3) are more close to Yak CSN2 and CSN3 gene respectively.

## Data availability

The data that support the findings of this study are available from the corresponding author upon reasonable request.

## References

1. Anand, V., Dogra, N., Singh, S., Kumar, S. N., Jena, M. K., Malakar, D., Dang, A. K., Mishra, B. P., Mukhopadhyay, T. K., Kaushik, J. K., & Mohanty, A. K. (2012). Establishment and characterization of a buffalo (Bubalus bubalis) mammary epithelial cell line. PloS one, 7(7), e40469. https://doi.org/10.1371/journal.pone.0040469.

2. Antonio Roberto Otaviano, Humberto Tonhati, Janete Aparecida Desidério Sena and Mario Fernando Cerón Muñoz (2005). Kappa-casein gene study with molecular markers in female buffaloes (Bubalus bubalis). Genetics and Molecular Biology, 28, 2, 237–241 (2005).

3. Azevedo, A. L., Nascimento, C. S., Steinberg, R. S., Carvalho, M. R., Peixoto, M. G., Teodoro, R. L., Verneque, R. S., Guimarães, S. E., & Machado, M. A. (2008). Genetic polymorphism of the kappa-casein gene in Brazilian cattle. Genetics and molecular research : GMR, 7(3), 623–630. https://doi.org/10.4238/vol7-3gmr428.

4. Bailey, T. L., Williams, N., Misleh, C., & Li, W. W. (2006). MEME: discovering and analyzing DNA and protein sequence motifs. Nucleic acids research, 34(Web Server issue), W369–W373. https://doi.org/10.1093/nar/gkl198.

5. Barłowska, J., Wolanciuk, A., Litwińczuk, Z., & Król, J. (2012). Milk Proteins’ Polymorphism in Various Species of Animals Associated with Milk Production Utility. http://dx.doi.org/10.5772/75168.

6. Batra, Kanisht & Nanda, Trilok & Kumar, Aman& Kumari, Rajni & Kumar, Vinay & Maan, Sushila. (2019). Molecular characterization of OAS1 as a biomarker molecule for early pregnancy diagnosis in Bubalus bubalis. Indian Journal of Biotechnology. 18. 97–107.

7. Blom, N., Gammeltoft, S., & Brunak, S. (1999). Sequence and structure-based prediction of eukaryotic protein phosphorylation sites. Journal of molecular biology, 294(5), 1351–1362. https://doi.org/10.1006/jmbi.1999.3310.

8. Blom, N., Sicheritz-Pontén, T., Gupta, R., Gammeltoft, S., & Brunak, S. (2004). Prediction of post-translational glycosylation and phosphorylation of proteins from the amino acid sequence. Proteomics, 4(6), 1633–1649. https://doi.org/10.1002/pmic.200300771.

9. Chakraborty, N., Besra, A., & Basak, J. (2020). Molecular Cloning of an Amino Acid Permease Gene and Structural Characterization of the Protein in Common Bean (Phaseolus vulgaris L.). Molecular biotechnology, 62(3), 210–217. https://doi.org/10.1007/s12033-020-00240-4.

10. Darshan Raj Gangaraj,_, Swathi Shetty, M.G. Govindaiah, C.S. Nagaraja, S.M. Byregowda, M.R. Jayashankar (2008). Molecular characterization of kappa-casein gene in buffaloes. ScienceAsia, doi: 10.2306/scienceasia1513-1874.2008.34.435.

11. Donald K. Layman, Bo Lo nnerdal, and John D. Fernstrom (2018). Applications for a-lactalbumin in human nutrition. Nutrition ReviewsVR Vol. 76(6):444–460. doi: 10.1093/nutrit/nuy004.

12. Elisabeth Gasteiger, Alexandre Gattiker, Christine Hoogland, Ivan Ivanyi, Ron D. Appel and Amos Bairoch (2003). ExPASy: the proteomics server for in-depth protein knowledge and analysis. Nucleic Acids Research, 2003, Vol. 31, No. 13 DOI: 10.1093/nar/gkg56.

13. Ertuğrul Filiz and Ibrahim Koc (2014). In silico sequence analysis and homology modeling of predicted beta-amylase 7-like protein in Brachypodium distachyon L.. J. BioSci. Biotech. 2014, 3(1): 61–67.

14. Geourjon, C., & Deléage, G. (1995). SOPMA: significant improvements in protein secondary structure prediction by consensus prediction from multiple alignments. Computer applications in the biosciences : CABIOS, 11(6), 681–684. https://doi.org/10.1093/bioinformatics/11.6.681.

15. Joanna Barłowska, Anna Wolanciuk, Zygmunt Litwińczuk and Jolanta Król (September 12th 2012). Milk Proteins’ Polymorphism in Various Species of Animals Associated with Milk Production Utility, Milk Protein, Walter L. Hurley, IntechOpen, DOI: 10.5772/50715.

16. John Bonsing, Jennifer M. Ring A, A. Francis StewartB and Antony G. Mackinlay (1988). Complete Nucleotide Sequence of the Bovine /3 -casein Gene. Aust. J. BioI. Sci., 1988, 41, 527–37.

17. Kishore, A., Mukesh, M., Sobti, R. C., Kataria, R. S., Mishra, B. P., & Sodhi, M. (2014). Analysis of genetic variations across regulatory and coding regions of kappa-casein gene of Indian native cattle (Bos indicus) and buffalo (Bubalus bubalis). Meta gene, 2, 769–781. https://doi.org/10.1016/j.mgene.2014.10.001.

18. Mark V. Berjanskii Æ David S. Wishart (2008). Application of the random coil index to studying protein flexibility. J Biomol NMR (2008) 40:31–48 DOI 10.1007/s10858-007-9208-0.

19. Sandeep Kumar, Chung-Jung Tsai, Ruth Nussinov (2000). Factors enhancing protein thermostability, Protein Engineering, Design and Selection, Volume 13, Issue 3, March 2000, Pages 179–191, https://doi.org/10.1093/protein/13.3.179.

20. Shental-Bechor, D., & Levy, Y. (2008). Effect of glycosylation on protein folding: a close look at thermodynamic stabilization. Proceedings of the National Academy of Sciences of the United States of America, 105(24), 8256–8261. https://doi.org/10.1073/pnas.0801340105.

21. Shi, S. P., Qiu, J. D., Sun, X. Y., Suo, S. B., Huang, S. Y., & Liang, R. P. (2012). PLMLA: prediction of lysine methylation and lysine acetylation by combining multiple features. Molecular bioSystems, 8(5), 1520–1527. https://doi.org/10.1039/c2mb05502c.

22. Timothy L. Bailey, Mikael Boden, Fabian A. Buske, Martin Frith, Charles E. Grant, Luca Clementi, Jingyuan Ren, Wilfred W. Li, William S. Noble (2009). MEME SUITE: tools for motif discovery and searching, Nucleic Acids Research, Volume 37, Issue suppl_2, 1 July 2009, Pages W202–W208, https://doi.org/10.1093/nar/gkp335.

23. Vinesh P.V., Biswajit Brahma, Rupinder Kaur, Tirtha Kumar Datta, Surender Lal Goswami, Sachinandan De (2013). Characterization of β-casein gene in Indian riverine buffalo. Gene. 2013 Sep 25;527(2):683–8. doi: 10.1016/j.gene.2013.06.029. Epub 2013 Jun 27.

24. Wayne S. Bawden, Robert J. Passey & Antony G. Mackinlay (1994) The Genes Encoding the Major Milk-Specific Proteins and Their Use in Transgenic Studies and Protein Engineering, Biotechnology and Genetic Engineering Reviews, 12:1, 89–138, DOI:10.1080/02648725.1994.10647910.

